## Supplementary information for "p23-FKBP51 chaperone complex regulates tau aggregation"

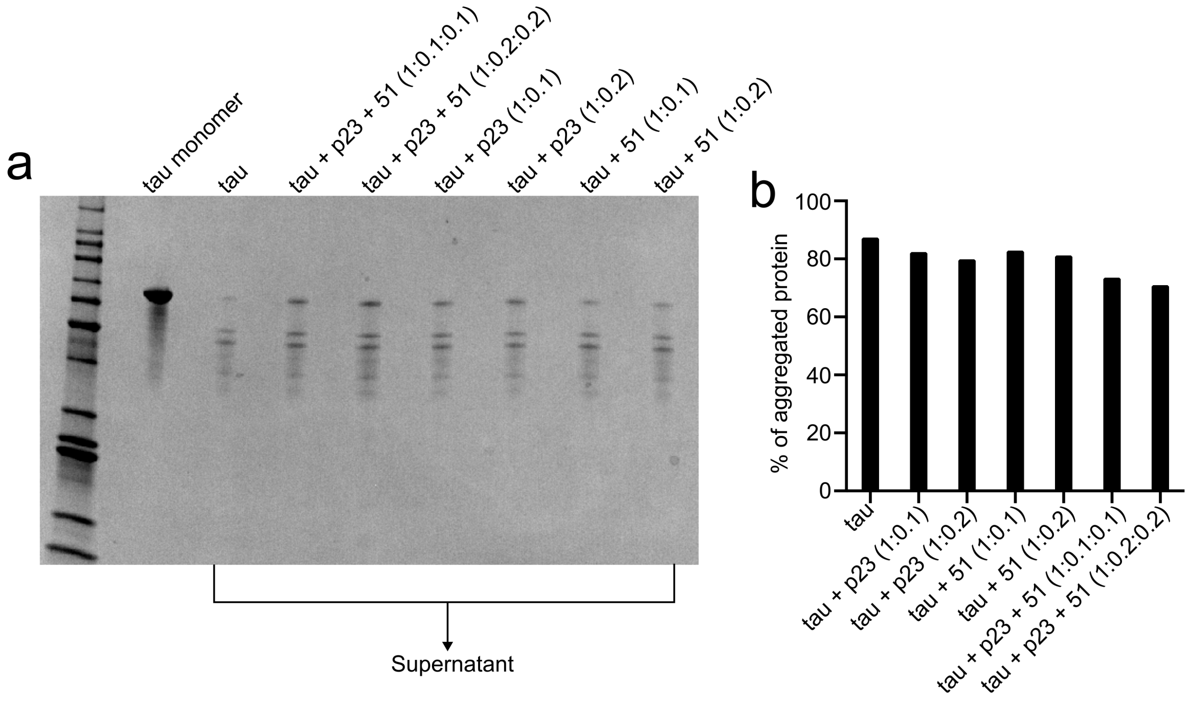


**Supplementary Fig. 1 | Determination of the amount of aggregated protein. a,** SDS-PAGE gel of tau monomer and supernatant (SN) (after pelleting down the fibrils) of tau either in the absence or presence of different co-chaperones. The fibril samples were collected after four days of aggregation. **b**, Calculation of the amount of aggregated protein. The amount of aggregated protein was calculated by comparing the intensity of the supernatant (SN) band to the tau monomer band.

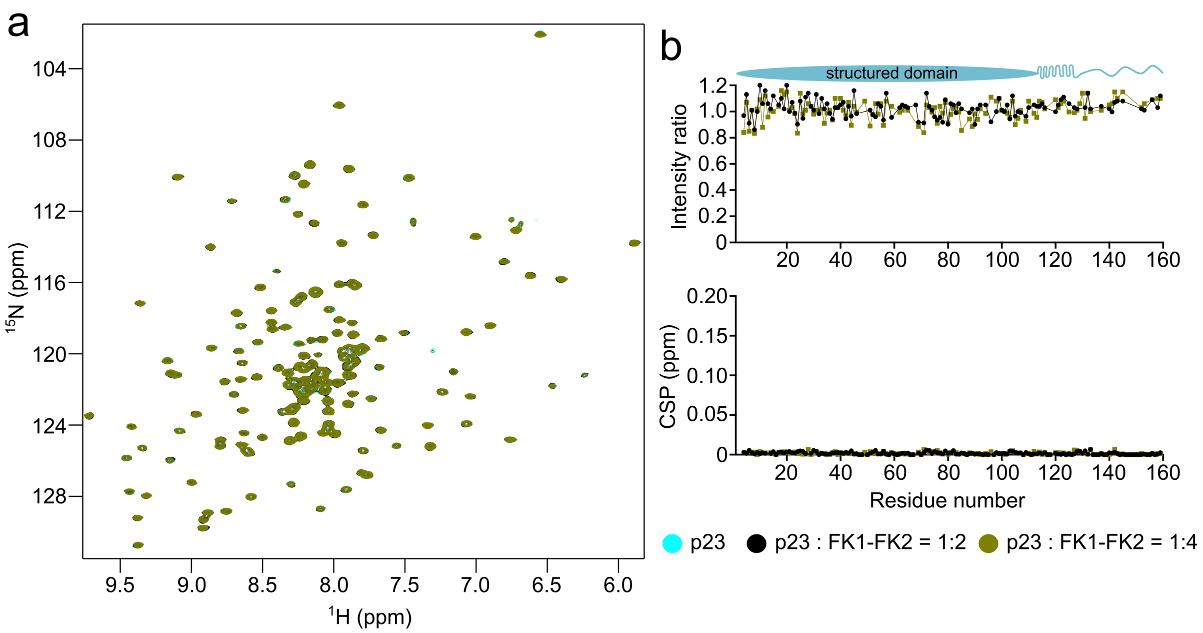


**Supplementary Fig. 2 | Interaction of p23 with FK1-FK2. a,** 2D $^{1}$H-$^{15}$N TROSY HSQC spectra of $^{15}$N-labeled p23 in the absence (cyan) or presence of two-fold (black) and four-fold (deep green) molar excess of unlabeled FK1-FK2. **b**, Changes in the intensities (top) and chemical shift perturbations (CSPs) (bottom) of the cross peaks in the TROSY-HSQC spectrum of p23 upon the addition of two-fold (black) and four-fold (deep green) molar excess of unlabeled FK1-FK2. The domain diagram of p23 is shown at the top.

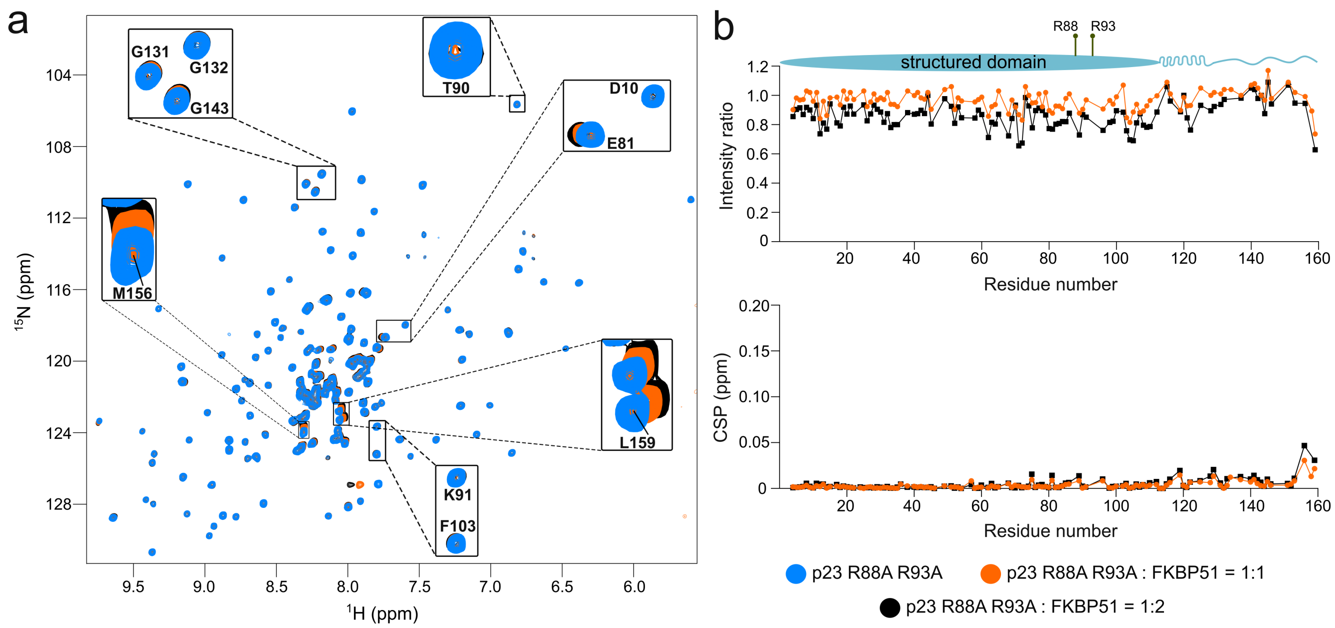


**Supplementary Fig. 3 | Interaction of R88A/R93A-mutant p23 with FKBP51. a,** 2D $^{1}H$-$^{15}N$ TROSY HSQC spectra of deuterated, $^{15}N$-labeled R88A/R93A-mutant p23 either in the absence (blue) or presence of equimolar (orange) and two-fold (black) molar excess of unlabeled FKBP51. **b**, Changes in the intensities (top) and chemical shift perturbations (CSPs) (bottom) of the cross peaks in the TROSY-HSQC spectrum of R88A/R93A-mutant p23 upon the addition of equimolar (orange) and two-fold (black) molar excess of FKBP51. The domain diagram of p23 with the position of R88 and R93 is shown above.

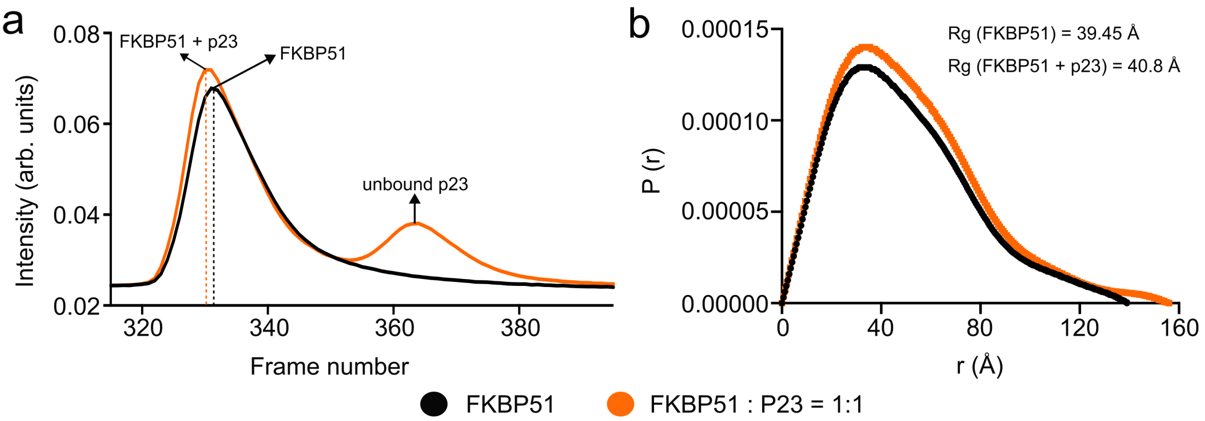


**Supplementary Fig. 4 | SEC-SAXS analysis of the p23-FKBP51 complex. a,** Elution profile of free FKBP51 (black) and the p23-FKBP51 complex (orange) from the size exclusion chromatography. **b**, P(r) distribution curve of the free FKBP51 (black) and the p23-FKBP51 complex (orange) calculated from SAXS. The values of the radius of gyration (Rg) of free FKBP51 and the p23-FKBP51 complex are indicated.

| | |
|---|---|
| HADDOCK score | -146.9 ± 3.4 |
| Cluster size | 92 |
| RMSD from the lowest-energy structure | 0.9 ± 0.5 |
| Van der Waals energy | -44.2 ± 5.7 |
| Electrostatic energy | -668.3 ± 41 |
| Desolvation energy | 27 ± 3.5 |
| Buried surface area | 2014.9 ± 115.2 |
| Z-score | -1.9 |

**Supplementary Table 1 | HADDOCK docking statistics of the structure of p23-FKBP51 complex**

| Mutant | Forward primer | Reverse primer | Template |
|---|---|---|---|
| p23 (1-119) | 5´CAGATGAAGACATGTCTAATTAAGATCG TTTCTCTGAGATGATG 3´ | 5´CATCATCTCAGAGAAACGATCTTAA TTAGACATGTCTTCATCTG 3´ | p23 |
| p23 R88A | 5´CAGTCATGGCCAGCGTTAACAAAAGAA AGG 3´ | 5´CCTTTCTTTTGTTAACGCTGGCCATG ACTG 3´ | p23 |
| p23 R88A R93A | 5' CCAGCGTTAACAAAAGAAGCGGCAAAGC TTAATTGGC 3' | 5' GCCAATTAAGCTTTGCCGCTTCTTTTG TTAACGCTGG 3' | p23 R88A |

**Supplementary Table 2│ Details of the primers to generate different mutants of p23**
